## Supplementary Information for "Molecular mechanisms of microbiome modulation by the eukaryotic secondary metabolite azelaic acid"

**Supplementary Figures**

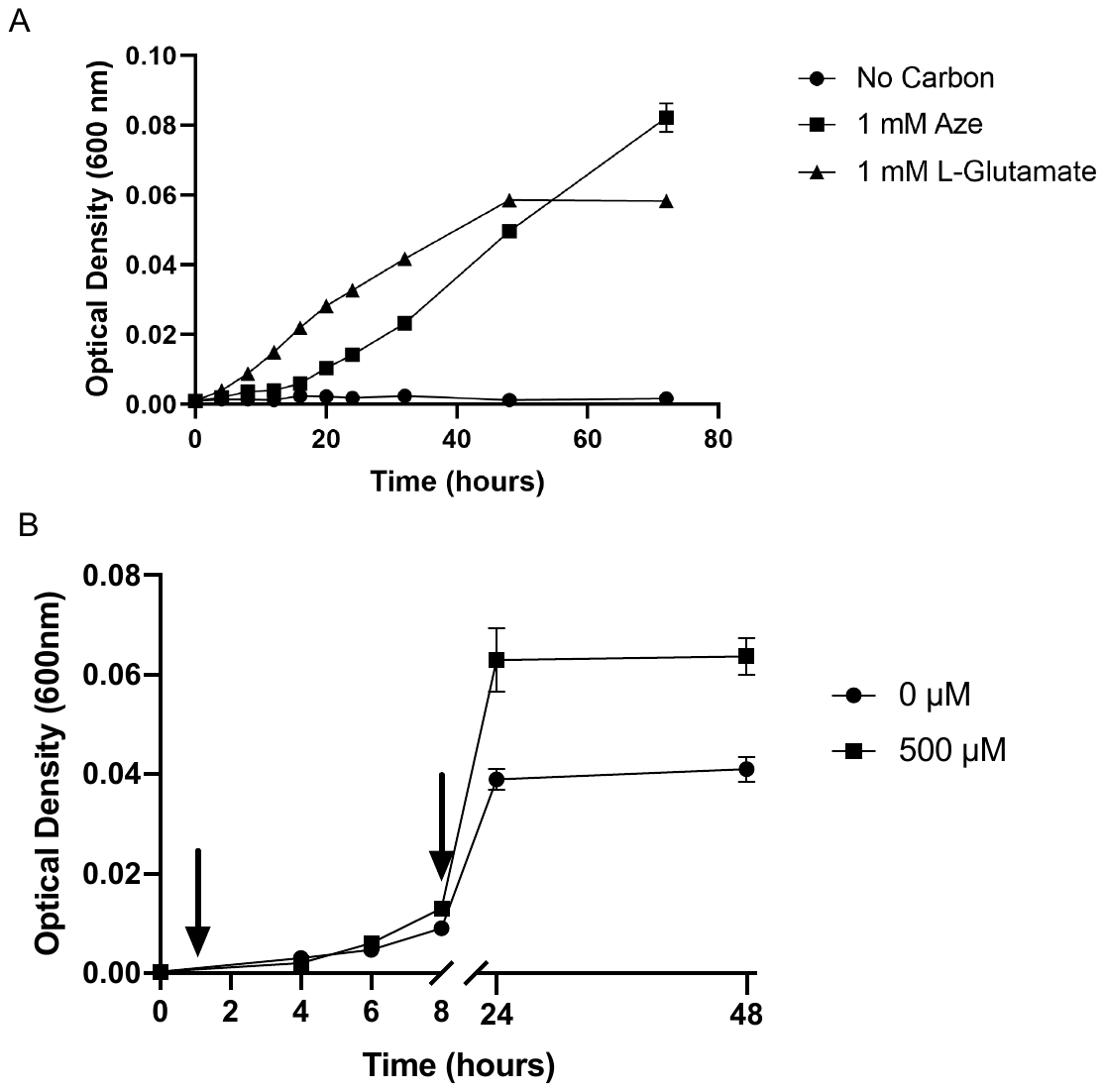

**Figure S1. Effect of Aze on growth of *Phycobacter*.** **a,** as a sole carbon source (1 mM) and **b**, in comparison to a control. For **(b)** Aze was added at T=0 hours to treatments while controls received an equivalent volume of Milli-Q water. RNA samples were collected at the times marked by arrows (T=0.5 hours and T=8 hours). Error bars represent standard deviation of triplicate cultures.

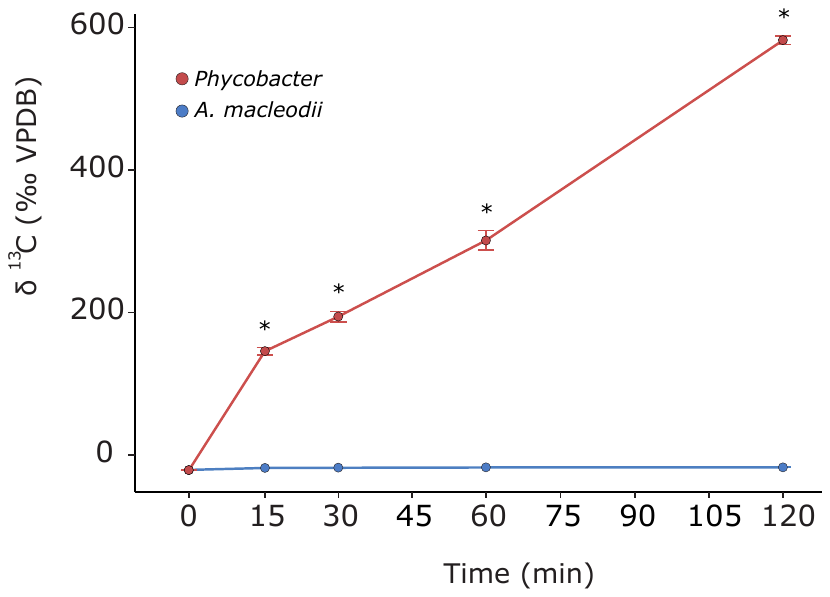

**Figure S2. ^13^C-Aze assimilation by *Phycobacter* and *A. macleodii* (reported as δ^13^C [‰, VPDB]) during a two-hour incubation.** Asterisks denote significant differences between the two bacteria (repeated measure ANOVA, *n*=3, *p*<0.001). Error bars indicate the standard deviation of the mean.

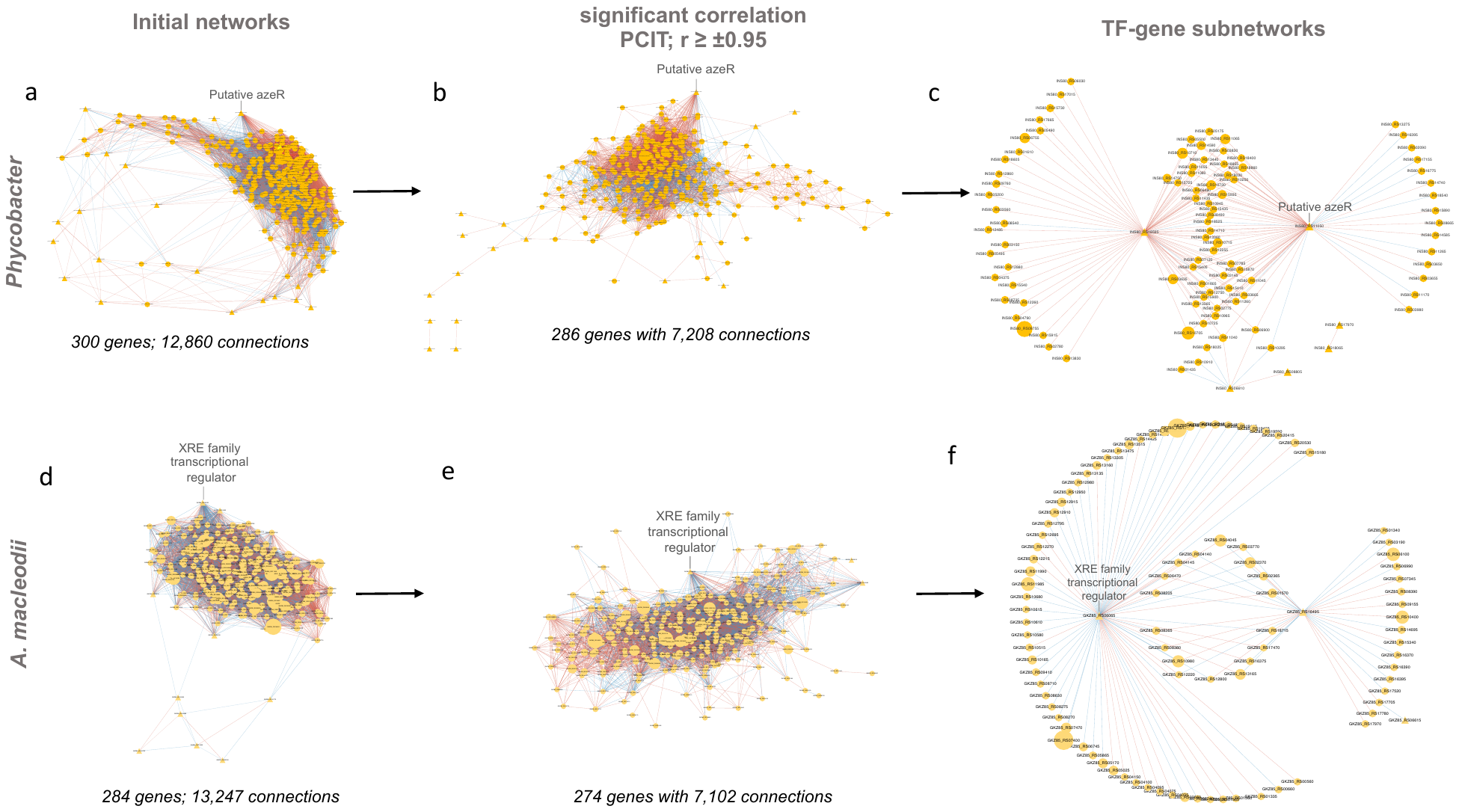

**Figure S3.** **Transcriptional coexpression networks constructed using the Partial Correlation coefficient with Information Theory algorithm in *Phycobacter* and *A. macleodii*.** Initial networks (**a, d**) consisted of key transcriptional factors (TFs) (identified by regulatory impact factor analysis) and differentially expressed genes*.* Nodes are depicted either as circles for genes or triangles for TFs. Edges represent significant interactions between nodes. Edge color represents directions of the interaction (red=positive correlation and blue=negative correlation). The size of the node corresponds to the normalized mean expression values in Aze-treated samples. Subnetworks were extracted from initial networks based on significant co-expression correlation (PCIT; r ≥ ±0.95) (**b, e**) and those containing only hub genes (identified based on RIF scores, differential expression, and the degree centrality) and their connected genes (**c, f**).

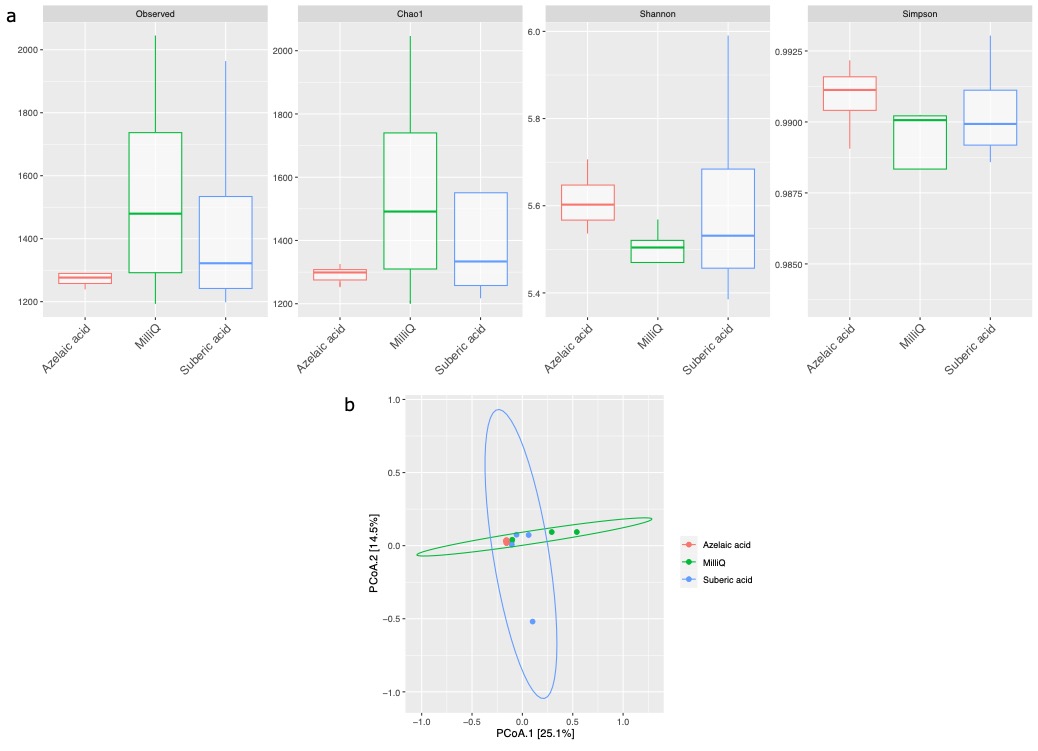

**Figure S4. Effect of Aze treatment on the alpha and beta-diversity of soil. a**, Alpha-diversity indices of observed OTUs, Chao1, Shannon and Simpson across the Aze-treated and control samples. **b**, PCoA of Bray-Curtis distances (PERMANOVA: *R*^2^ = 0.73; *p*<0.001) between samples. The two principal components (PCoA1 and PCoA2) explained 25.1% and 14.5% variance, respectively.

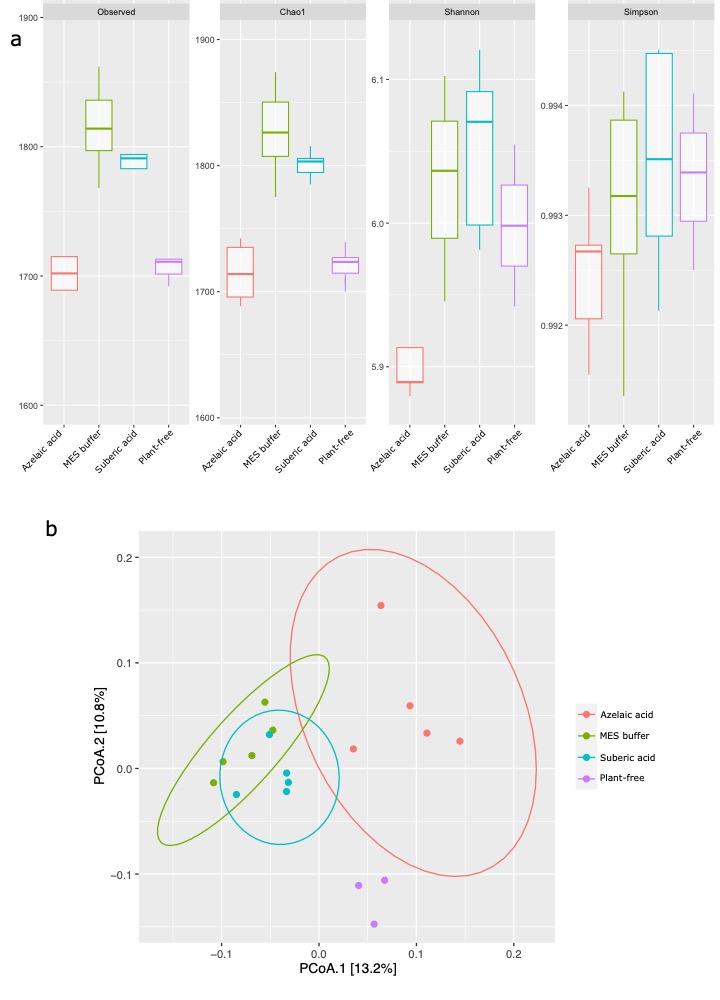

**Figure S5. Effect of Aze treatment on the alpha and beta-diversity of *A. thaliana* root microbiome. a**, Alpha-diversity indices of observed OTUs, Chao1, Shannon and Simpson across the Aze-treated and control treatments. **b**, PCoA of Unweighted Unifrac distances between Aze-treatment and controls (PERMANOVA, *p*<0.05). The two principal components (PCoA1 and PCoA2) explained 13.2% and 10.8% variance, respectively.

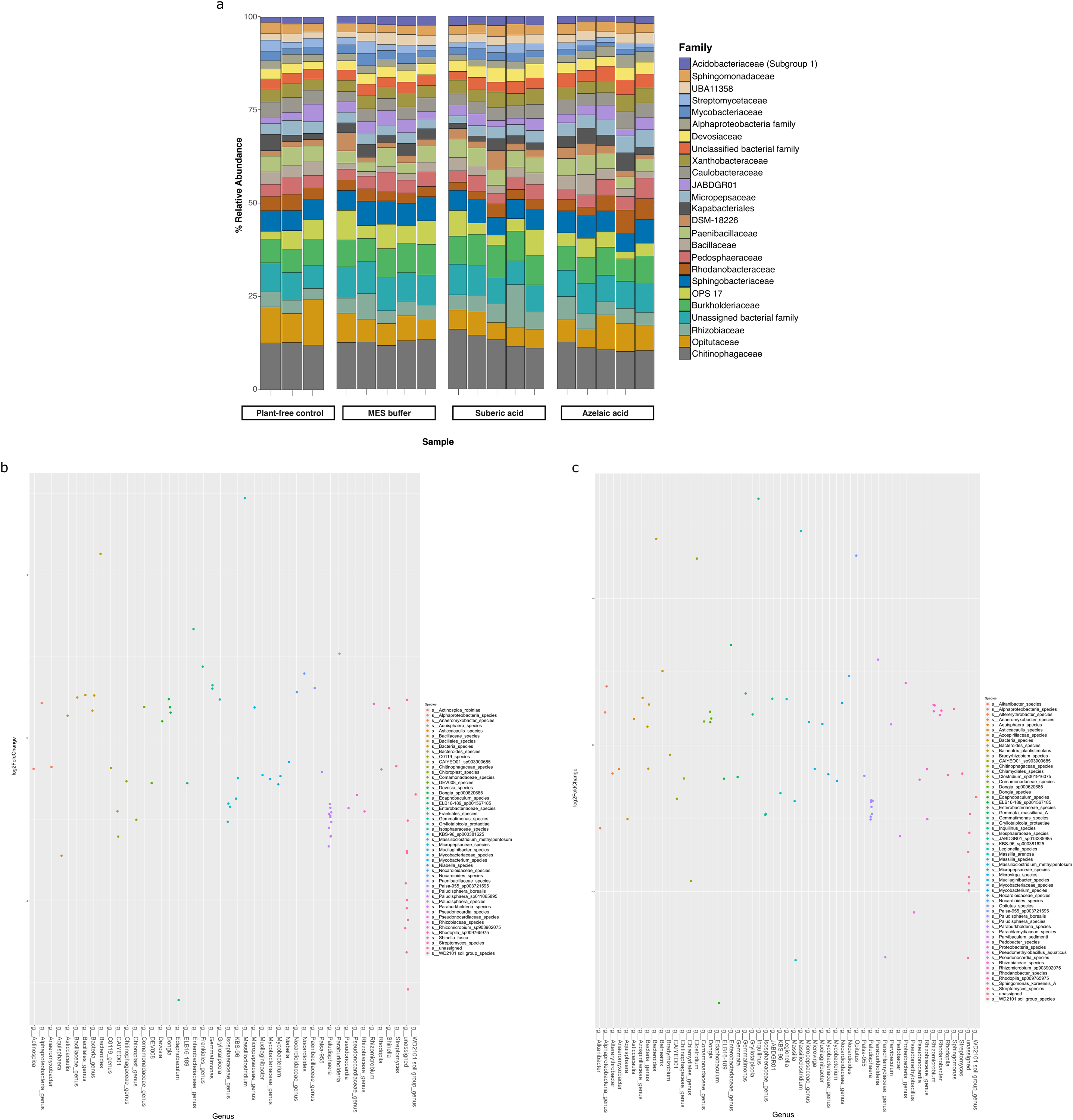

**Figure S6. Bacterial diversity in the roots of *A. thaliana* treated with Aze. a,** Relative abundance of the top 25 microbial families based on 16S rRNA gene amplicon sequencing of *A. thaliana* root-microbiome following Aze infiltration. Plant-free control is untreated soil and MES buffer and suberic acid are treatment controls. **b,** Distribution of the ASVs belonging to significantly differentially abundant taxa between the Aze-treated and MES buffer samples according to their log_2_ fold-change and *p*-adjusted values. The bubble color denotes the species belonging to each genus on the x-axis. **c,** Distribution of the ASVs belonging to significantly differentially abundant taxa between the Aze-treated and suberic acid samples according to their log_2_ fold-change and *p*-adjusted values. Bubble colors denote the species belonging to each genus on the x-axis.
